# A consistent evaluation of miRNA-disease association prediction models

**DOI:** 10.1101/2020.05.04.075754

**Authors:** Ngan Thi Dong, Megha Khosla

**Affiliations:** L3S Research Center, Leibniz University Hannover, Hannover, 30167, Germany

## Abstract

**Motivation:** A variety of machine learning based approaches have been applied to predicting miRNA-disease association. Although promising, the evaluation set up to measure prediction performance is inconsistent making it difficult to assess the actual progress. A more acute problem is that most of the models overlook the problem of data leakage due to the use of precomputed miRNA and disease similarity features.

**Results:** We unearth a crucial problem of data leakage in evaluation of machine learning models for miRNA-disease association prediction. In particular, information from test set, in the form of precomputed input features for miRNA and disease, is used during training of the model. Moreover, we point out problems in the widely used performance metrics used in model evaluation. While resolving the issues of data leakage and model evaluation, we perform an indepth study of 3 recent models along with our proposed 9 variants of these models. Our proposed variants have resulted in improvements in Average Precision scores (as compared to original models) by approximately 287.7% and 36.7% on HMDDv2.0 (AP:0.504) and HMDDv3.0 (AP: 0.216) datasets respectively.

**Availability and Implementation:** We release a unified evaluation framework including all models and datasets at https://git.l3s.uni-hannover.de/dong/simplifying_mirna_disease.

## 1 Introduction

MicroRNA or miRNA is a class of noncoding RNA with a length of approximately 22 nucleotides that is involved in the regulation of gene expression. miRNA can up-regulate or down-regulate genes and thus indirectly affect the transcription process. It has been observed that many of the miRNAs are associated with the development of complex diseases in humans. Identification of potential association between miRNA and disease would therefore help in clinical diagnosis, treatment and in finding potential drug targets.

As the biological methods for detecting these associations are expensive and time-consuming, recent years have seen an upsurge in computational approaches for predicting miRNA-disease associations. These approaches can be broadly grouped into three classes: scoring based methods, network topology based methods and machine learning based methods. Assuming that the miRNA pairs linked to common diseases are functionally more related [16, 14] proposed scoring systems to prioritize miRNA-disease associations. A more sophisticated scoring scheme while integrating information from miRNA and disease similarity networks was proposed in [19]. The second class of methods, i.e the network based algorithms [10, 4] construct miRNAs and/or disease similarity networks and aims at efficient transferring known miRNA-disease association labels between similar miRNAs and/or similar diseases in the network. Chen et al. [2] employ repeated random walks with restarts over the miRNA functional similarity network and prioritize candidate miRNA-disease associations using the final stable walk probability. In this work we mainly focus on machine learning based methods which typically build classification or regression models on input features extracted from different classes of heterogeneous information like disease semantic information, miRNA–disease association information and miRNA function information and sometimes also miRNA sequence information. For example, RBMMMDA [3] uses Restricted Boltzmann machine for the prediction task. DRMDA[1] applies a support vector machine model over the miRNA and disease features extracted using stacked autoencoders with unsupervised pre-training. DeepMDA[6] is a deep ensemble model which extracts features using stacked autoencoders and predicts finals scores of associations using a 3-layer neural network. A comparative study of various computational methods for miRNA-disease association prediction can be found in [**?**].

More recent works (published in January 2020) include Epmda [5] which first extracts edge perturbation based features from the miRNA and disease similarity networks and then train a multilayer perceptron regression model to prioritize miRNA-disease associations. Another work, Dbmda [20] uses autoencoders to extract latent representations of miRNA and disease similarity as well as miRNA sequence similarity to further train a rotation forest classifier to predict prospective associations. Nimgcn [11] uses the graph convolution networks (GCNs)[8] to extract representations of miRNA and similarity networks and further use them to train a neural matrix completion network. All of these methods integrate different heterogeneous data sources, most notably, miRNA functional similarity, disease similarity extracted from the disease ontology, known miRNA-disease associations and in some cases also miRNA sequence similarity.

### Problem of Data Leakage

While information integration from multiple sources has shown to improve performance, in this paper we show that the evaluation schemes in such a multi-source scenario suffers from a data leakage problem. Data leakage in machine learning refers to scenario when information from the test set, that should in principle be inaccessible during training, is used to create the model. To understand the problem of data leakage in miRNA-disease association predictive models, consider the concrete example of using precomputed miRNA functional similarities during model training. miRNA functional similarity (for details see Section 2) is computed using the known miRNA-disease associations and disease semantic information. More often, the precomputed miRNA functional similarity from MISIM database is used in model building without giving proper attention to the actual train/test split giving rise to data leakage. This implies that some of the associations which are to be tested by the model might be present in the association network that were used to compute the similarity features for example miRNA functional similarity. This leads to erroneous evaluation of models and overestimation of actual learning/prediction capabilities of the models. As an example, [**?**] also observes a fall in the performance of various models when trained on HMDD v2.0 dataset and tested on HMDD v3.0 dataset which the authors attributed to overfitting of models when trained and tested on the same dataset. We clarify this issue by showing that this discrepancy occured because of the data leakage due to the use of precomputed miRNA functional similarities deposited in databases and in some cases the use of the entire association networks to compute for example Gaussian Interaction Profile kernels.

### Limitations of the evaluation setup

In addition, there has been a huge diversity in the experimental datasets used and evaluation metrics. Some models only report Area under the Receiver Operating Characteristic Curve (AUROC) scores which is typically more optimistic when the size of positive (actual associations) class is much smaller than the negative (non associations) class. A better performance metric is Area under Precision-Recall curve (AUPRC) which takes into account both precision and recall of the model at different decision thresholds. As computing AUPRC requires interpolating between adjacent supporting points, it too overestimates the actual performance in extreme cases, for example when there are just two pints in precision-recall curve. In this work, we instead use Average Precision (AP) (more details in Section 4.2) which summarizes a precision-recall curve as the weighted mean of precisions achieved at each threshold, with the increase in recall from the previous threshold used as the weight. It does not use linear interpolation and provides a more reliable performance estimate than AUPRC.

### Our Contributions

In this work, we perform an indepth study of 3 recent machine learning models, namely Epmda, Dbmda and Nimgcn. In addition to resolving the problems of data leakage and evaluation setup, we propose and study 9 variants of the studied models. Through a comparative analysis of resulting 12 models over two widely used datasets our work offers deeper insights into the workings of very recent machine learning models for the miRNA-disease association prediction task. To summarize, we make the following contributions.

- We unearth a crucial problem of data leakage in evaluation of machine learning models for miRNA-disease association prediction. We develop and release an integrated evaluation framework in which various miRNA and disease similarities are computed for each training set thus disallowing any data leakage from the test set.
- We thoroughly investigate 3 state of the art models including 9 of their simpler variants showing advantages of reduced model complexities. Our proposed simpler variants have resulted in one of the best Average Precision score observed so far on HMDD v2.0 [12] and HMDD v3.0 [7] datasets.
- We compared the effect of using different input miRNA and disease similarity features and find that using Gaussian Interaction Profile (GIP) kernel similarity leads to better performance.

## 2 Preliminaries

In this paper, we study supervised machine learning methods for the miRNA-disease associations prediction problem. That is to say, from a set of known miRNA-disease associations, learn a model to predict potential miRNA-disease associated pairs. Our studied models treat that problem as either a binary classification task where each pair is represented as a feature vector fed as input to a classifier, or as a matrix completion problem. In order to construct the feature vector or build the matrix, different similarities measures have been used. This section will give a brief overview of all the similarity measures employed by the models studied in this work.

### Disease Similarity

Disease Semantic Similarity[16] computes the semantic similarity of two diseases based on their relative positions on the disease directed acyclic graph (DAG) which is derived from the MeSH^1^ disease descriptor database. Specifically in the disease DAG the nodes represent diseases and directed links from parent nodes to child nodes encode a ‘is-a’ relationship. The contribution of disease *A* to the semantic value of disease *t* is then defined as:

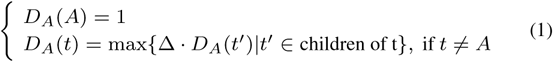

where Δ is the decaying factor and is set to 0.5 as in [16]. Then the semantic value for disease *A* is calculated as 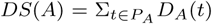, where *P*_*A*_ is the set of *t*’s ancestors. Finally the **semantic similarity** between two diseases *A* and *B* is given by:

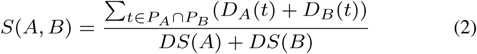

where *P*_*A*_ and *P*_*B*_ are the set of ancestors for diseases *A* and *B* respectively. However, Xuan et al. [17] argue that the more a particular disease appears as an ancestor of other diseases the less specific it is for a particular disease. Given that the likelihood of a disease node *t* appearing as ancestors of all other diseases, [17] quantifies the information content of *t, IC*(*t*), as the negative log of the likelihood, i.e. *IC*(*t*) = − log(*p*(*t*)). The phenotype similarity score between two diseases is then given by

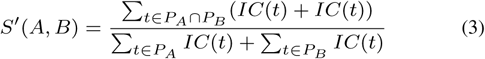

The updated similarity score between two diseases is then calculated as an average of their semantic similarity and phenotype similarity scores.

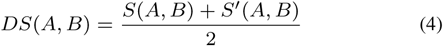

For simplicity, from now on, without explicitly given the formula, by saying **disease semantic similarity**, we mean the disease semantic similarity given in Equation (4).

### miRNA functional similarity [16]

The miRNA functional similarity is estimated based on the similarities between the diseases to which they are associated. In particular, first the similarity between disease *d*_*i*_ and a group of disease *DG* is computed as 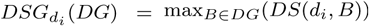. Given that 𝒟_*i*_ denote the set of disease associated with miRNA *m*_*i*_, the functional similarity between two miRNAs *m*_1_, *m*_2_ is then calculated as:

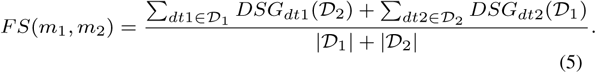

### miRNA sequence similarity [11]

The miRNA sequence is first mapped to its 2 dimensional (2D) representation where each nucleotide in its sequence is converted to a 2-dimensional vector. For *i*th nucleotide in a particular miRNA sequence, its 2D encoding is computed by the recursive relation *nt*_*i*_ = *nt*_*i*−1_*θ*(*nt*_*i*−1_ − *λ*_*i*_) such that *nt*_0_ = (0.5, 0.5) and the decaying factor *θ* = −0.5. The parameter *λ*_*i*_ is set as follows

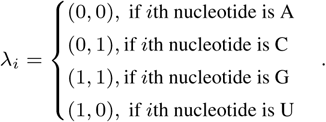

The sequence similarity of two miRNAs *m*_*i*_ and *m*_*j*_ is then defined as the Euclidean distance between their two vectors in the above defined 2D space.

### Gaussian Interaction Profile (GIP) Kernel Similarity [15]

Given the miRNA-disease association matrix *MD*, the GIP kernel similarity between two miRNAs is defined as:

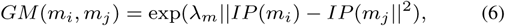

where *IP* (*m*_*i*_) and *IP* (*m*_*j*_) correspond to the *i*th and *j*th rows of matrix *MD* respectively. The parameter *λ*_*m*_ is set to.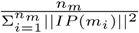 where *n*_*m*_ is the total number of miRNAs. The Gaussian Interaction Profile kernel similarity between two diseases is calculated similarly as:

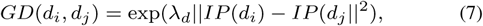

where *IP* (*d*_*i*_) and *IP* (*d*_*j*_) correspond to the *i*th and *j*th column of matrix *MD* accordingly. The parameter *λ*_*d*_ is set to 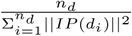 where *n*_*d*_ is the total number of diseases in *MD*.

## 3 Compared Models

In this section we describe the 3 recent models studied in this work along with our proposed variants. Our proposals for model variants are based on (i) reducing redundant non-linearities hence decreasing the number of model parameters (ii) identifying pitfalls in training data set for example the use of imbalanced training set in Epmda. In the following we provide details of the studied models and our proposed variants.

### Epmda [5]

The edge perturbation based method for miRNA-disease association prediction (Epmda) provides a new feature extraction method based on edge perturbation in miRNA-disease heterogenous network.

The heterogenous network *G* input to Epmda can be divided into three parts as follows:

- *The miRNA-disease bipartite network* consists of two type of nodes: one for miRNA and one for disease. There is no connection between two nodes of the same type and a link between a miRNA and a disease node indicates a known association.
- *The miRNA similarity network* is a weighted network where each node is a miRNA and the edge weight is equivalent to the `GIP` kernel similarity between two miRNA nodes (as defined in Equation (6)).
- *The disease similarity network* is a weighted network where each node represents one disease and the weight of the edges are the disease `GIP` kernel similarities (as given in Equation (7)).

The weighted adjacency matrix (*A*) of *G* is given as

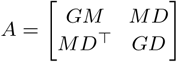

The problem of miRNA-disease association prediction is treated as the problem of predicting links between miRNA nodes and disease nodes on *G*. Feature extraction corresponding to edges in *G* is then proposed based on the network topology where each feature *f*_*i*_ reflects the disturbance level resulted from the deletion of that particular edge to the set of cycles (in which it participates) of length *i* in *G*. The higher the disturbance, the more important that particular edge is. A 5-layer Multilayer Perceptron Regression model is then trained using the extracted features with regularized square loss to predict whether there is a link between the miRNA-disease pair.

### Variants of Epmda

We investigated a variant of Epmda where we replaced the MLP regressor with a linear regressor. In addition, as elaborated later in Section 4.3.1, Epmda uses an imbalanced training set which might downgrade its performance. Therefore, besides the original proposed model, we also propose and investigate three variants of Epmda as summarized in Table 1. We call a data set that has equal number of positive and negative samples a balanced dataset. Epmda denotes the original proposed model which uses an imbalanced training dataset and the MLP regressor. Epmda1 is the model retrieved when we replace the MLP regression model with its linear counterpart. Epmda2 is obtained when we use a balance training set instead of the imbalanced one. Finally, Epmda3 combines the changes we made in Epmda1 and Epmda2. Namely, we both replace the MLP regression model with Linear regression model and use a balance training set to train the model.

**Table 1.** EPMDA and its variants

| Model | Training dataset |  | Regression Model |  |
| --- | --- | --- | --- | --- |
|  | IMBALANCED | BALANCED | MLP | LINEAR |
| EPMDA | ✓ | x | ✓ | x |
| EPMDA1 | ✓ | x | x | ✓ |
| EPMDA2 | x | ✓ | ✓ | x |
| EPMDA3 | x | ✓ | x | ✓ |

### Dbmda [20]

The distance-based sequence similarity for miRNA-disease association prediction (Dbmda) combines the information of miRNA sequence, miRNA functional similarity, miRNA-disease associations and disease semantic similarity. Dbmda architecture is illustrated in Figure 1. In particular for a given miRNA-disease pair it first concatenates the miRNA functional similarity and disease similarity features (computed in Section 2) and extracts a joint low dimensional representation using an autoencoder (denoted as Autoencoder 1 in Table 2). Also corresponding to miRNA, the sequence similarity features as described in Section 2 are fed to another autoencoder (denoted as Autoencoder 2 in table 2) to extract latent representation corresponding to miRNA sequence. The two newly extracted representations are concatenated and then fed to a rotation forest classifier to predict potential miRNA-disease associations.

**Table 2.** DBMDA and its variants

| Model | Autoencoder 1 | Autoencoder 2 | miRNA sequence features |
| --- | --- | --- | --- |
| DBMDA | ✓ | ✓ | ✓ |
| DBMDA1 | x | x | ✓ |
| DBMDA2 | ✓ | ✓ | x |
| DBMDA3 | x | x | x |

**Fig. 1.**
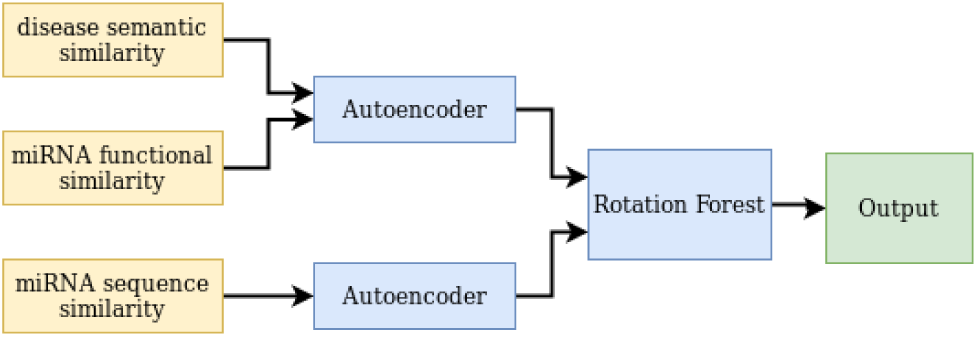
DBMDA model architecture

### Variants of Dbmda

We want to investigate the effectiveness of miRNA sequence similarity, autoencoders on Dbmda performance. Therefore, in this paper, apart from the original proposed model, we also report results from its three variants in which we either remove the auto-encoder, the miRNA sequence similarity features or both. As shown in Table 2, the Dbmda model corresponds to the original model. We obtain Dbmda1 from the removing the two autoencoders which are used to extract low dimensional representations from miRNA-disease pair and miRNA sequence similarity. Instead, for a particular miRNA-disease pair, we directly concatenate the miRNA functional, disease and miRNA sequence similarity to form a features vector. Dbmda2 corresponds to not using or removing the miRNA sequence similarity features from the input. For Dbmda3 we do not use miRNA sequence similarity features and any of the autoencoders. The concatenated features of miRNA-disease pair are directly used as input to the rotation forest classifier.

### Nimgcn [11]

Nimgcn operates on the heterogeneous miRNA-disease association network in which there are two type of nodes: miRNAs and diseases. miRNAs nodes form a miRNA functional similarity network where the edge weights correspond to the miRNA functional similarity scores as given in (5). Disease nodes form a disease semantic network where the edge weights are given by disease semantic similarity score calculated according to (4). A link between a miRNA and a disease node indicate a known association. Node representations are learnt using two Graph Convolution Networks (denoted as GCN encoders in figure 2) each for miRNA functional similarity network and disease similarity network separately. Non linear transformation or neural projections are applied over these learnt representations to extract latent features corresponding to miRNA and disease respectively. The predictions for miRNA-disease association ratings are then computed as the inner products of the latent features corresponding to miRNA and disease. Nimgcn architecture is depicted in Figure 2.

**Fig. 2.**
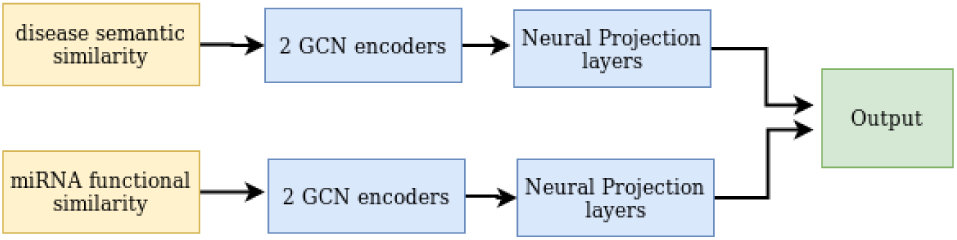
NIMGCN model architecture.

### Variants of Nimgcn

In this paper, apart from the original Nimgcn model, we also construct experiments on three of its variations in which we remove some part of the original model to make a simpler one. Details about those variants are given in table 3. For Nimgcn1, we remove the GCN encoders. miRNA functional and disease similarity scores are fed directly as input to the neural projections to learn hidden representation. For Nimgcn2, we remove the stack of three neural projections. Output from GCN encoders are used directly as miRNA and disease hidden representation. And for Nimgcn3, we use only one GCN encoder and one neural projection to learn the latent features of miRNA/disease.

**Table 3.** NIMGCN and its variants

| Model | Architecture |
| --- | --- |
| NIMGCN | original |
| NIMGCN1 | without GCNs |
| NIMGCN2 | 2 GCN layers, without neural projection |
| NIMGCN3 | 1 GCN layer and 1 neural projection layer |

## 4 Data and Experimental setup

### 4.1 Data Collection

We use two datasets derived from HMDD 2.0 [12] and HMDD 3.0 [7] as our evaluation datasets. The first dataset, which from now on we denoted as Hmdd2 consists of 5430 associations between 495 and 383 diseases. The second dataset, which from now on we denoted as Hmdd3, consists of 35362 associations between 1062 and 893 diseases.

The MESH terms for disease semantic similarity calculation are downloaded from the National Library of Medicine^2^. Diseases are matched by names. There are 55 and 360 disease names which are not found in MESH in Hmdd2 and Hmdd3 respectively. For those diseases, we fill in the similarity matrix with their corresponding Gaussian Interaction Profile (`GIP`) kernel similarity (calculated from the miRNA-disease association network) as described in Section 2. All miRNA sequence information was retrieved from miRBase [9]. The two datasets statistics are given in Table 4.

**Table 4.** The evaluated datasets statistics. *n*_*m*_, *n*_*d*_, *N* denote the number of miRNAs, diseases, and known associations respectively. *n*_*d*∉*MESH*_ refers to the number of diseases that are not found in MESH.

| Dataset | $n_m$ | $n_d$ | $N$ | $n_{d \notin MESH}$ |
| --- | --- | --- | --- | --- |
| HMDD2 | 495 | 383 | 5430 | 55 |
| HMDD3 | 1062 | 893 | 35362 | 360 |

### 4.2 Evaluation metrics

In terms of measurements, most of the paper in the fields can use either top K Predictive Rate, Area under the Receiver Operating Characteristic (AUROC) or Area under the Precision Recall Curve (AUPRC). However, for top K Predictive Rate, different value of K leads to different evaluations and in real applications, it is hard to specify the appropriate value of K. Therefore, we believe that using top K Predictive Rate is not recommendable. For AUROC score, as previously discussed in [18], AUROC score sometimes misleading for highly imbalanced datasets like our network of interest (the rate of positive link: negative link is 5430:184155 ≈ 0.0295 and 35362:948366 ≈ 0.0373 for Hmdd2 and Hmdd3 respectively). We argue that the use of AUPRC is more appropriate in this case. However, a point to be noted here is the calculation of the area under the precision-recall curve. The sklearn function to calculate the area under the cuve^3^ might suffer from the overestimation of the actual area because of linear interpolation. For example if there are just two points of x- and y-axis it computes the area under the line joining these two points which is clearly incorrect. We instead use Average Precision(AP)^4^ which summarizes a precision-recall curve as the weighted mean of precision achieved at each threshold (the weight here is the increase in recall from the previous threshold). AP is computed as:

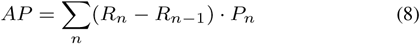

where *R*_*n*_, *P*_*n*_ are the recall and precision at the *n*th threshold correspondingly. Since Epmda and Nimgcn report AUROC as their evaluation metrics, for completeness, we report both AUROC and AP in our results.

### 4.3 Experimental setup

For all the models on a particular data set, we run 5 times 5-fold CV on the known miRNA-disease association set with different random seeds. Overall each model was trained/tested 25 times corresponding to 25 possible combination of training/testing splits. For simplicity, we call a training/testing split as a data split. More details about the data preparation process is given in Section 4.3.1. For all models, we use the same set of data splits.

In terms of the choice of similarity measure fed as input to the models, we experimented with three different type of similarities (input features): (i) the **`GIP`**: where we use the `GIP` kernel similarity for miRNA and disease; (ii) the `Functional+Semantic` where we use the miRNA functional similarity and disease semantic similarity given in Equation (4); and (iii) the `miRNA sequence+Semantic` where we use the miRNA sequence similarity and disease semantic similarity given in Equation (4).

We also construct experiments where we use the disease semantic similarity as stated in (2) instead of the one given in (4). However, from the results retrieved, we saw no significant difference between the two. Because of that reason and also due to space limit, we do not present the results corresponding to this experiment in this paper but details are available upon request.

#### 4.3.1 Training-testing data set up

##### Balanced training set

To create training splits we first randomly split all the known associations (positive examples) into 5 folds. The 4 folds constitute the training data corresponding to positive examples. To that we add an equal number of random sampled pairs which are not in those 4 folds to form the negative examples of the training set.

##### Test set preparation

To build the test set we use the remaining fold of the positive examples and mark all the others possible miRNA-disease combination which are not in the set of known association as negative examples. In our evaluation set up we use all negative pairs in the test set as opposed to artificially creating a balanced test set by sampling a smaller number of negative examples. This setting is more realistic as in real time the task will be to rank the small number of prospective associations among a large number of possible associations. Such a setting is also emphasized in [18].

A different train/test set up is used in the original papers of Nimgcn and Dbmda which explicitly mark all unknown pairs as negative samples and divide the set of all possible pairs into 5 folds. Each time, a fold is used for testing and the others for training. For example, in case of Hmdd2 dataset, the 5430 known miRNA-disease associations are used as positive samples while the other 184155 - 5430 = 183725 pairs are explicitly marked as negative samples. Instead of a complete test set of more than 180k pairs, their test set size is just around 36k pairs. There are two problems with this kind of configuration: (i) their training set is highly biased as the number of negative examples are much more than that of positive examples which is often detrimental to the model behavior, for example training error might be close to zero if the model labels all examples as negative. (ii) the test set contains much smaller amount of negative examples than the actual, hence the model is not tested in a realistic setting where it is required to predict/prioritize a small number of associations from a large number of possible combinations.

For Epmda, the authors also divide only the positive associations into 5 folds like our setting. However, for each data split, their training data composed of 4 folds of positive samples and all the other possible pairs as negative samples which also results in an imbalanced training set. We test the Epmda model using their original setting where they use an imbalanced training set (referred to as Epmda in in Table 1, Figures 3 and 4) and in our setting where a balanced training dataset is used (referred to as Epmda3 in tables 1, 7 and Figure 4). We see a considerable improvement in the model performance on Hmdd2 dataset when a balanced training set was used.

**Table 5.** Mean AUROC and AP scores (with standard deviation) for Nimgcn and its variants on HMDD2 and HMDD3 datasets. The original NIMGCN model uses Functional+Semantic similarity.

|  | Method | GIP |  | Functional+Semantic |  | miRNA sequence+Semantic |  |
| --- | --- | --- | --- | --- | --- | --- | --- |
|  |  | AUROC | AP | AUROC | AP | AUROC | AP |
| HMDD2 | NIMGCN | 0.906 ± 0.006 | 0.152 ± 0.009 | 0.857 ± 0.072 | **0.164 ± 0.032** | 0.850 ± 0.005 | 0.056 ± 0.003 |
|  | NIMGCN1 | 0.913 ± 0.004 | 0.146 ± 0.010 | 0.889 ± 0.008 | 0.161 ± 0.011 | 0.848 ± 0.006 | 0.056 ± 0.003 |
|  | NIMGCN2 | 0.889 ± 0.017 | **0.174 ± 0.015** | 0.856 ± 0.008 | 0.148 ± 0.014 | 0.892 ± 0.008 | **0.177 ± 0.011** |
|  | NIMGCN3 | 0.779 ± 0.019 | 0.086 ± 0.010 | 0.849 ± 0.007 | 0.149 ± 0.014 | 0.883 ± 0.006 | 0.166 ± 0.012 |
| HMDD3 | NIMGCN | 0.934 ± 0.007 | 0.199 ± 0.006 | 0.887 ± 0.086 | 0.158 ± 0.045 | 0.856 ± 0.006 | 0.036 ± 0.003 |
|  | NIMGCN1 | 0.939 ± 0.002 | 0.205 ± 0.007 | 0.858 ± 0.043 | 0.073 ± 0.025 | 0.858 ± 0.003 | 0.038 ± 0.001 |
|  | NIMGCN2 | 0.918 ± 0.024 | **0.212 ± 0.013** | 0.872 ± 0.022 | 0.162 ± 0.011 | 0.813 ± 0.067 | 0.070 ± 0.042 |
|  | NIMGCN3 | 0.913 ± 0.004 | 0.126 ± 0.006 | 0.913 ± 0.004 | **0.216 ± 0.006** | 0.909 ± 0.004 | **0.171 ± 0.006** |

**Table 6.** Mean AUROC and AP scores (with standard deviation) for Dbmda and its variants on HMDD2 and HMDD3 datasets. The original Dbmda model uses Functional+Semantic similarity along with miRNA sequence similarity. Results for Dbmda and Dbmda1 are not presented in the last column as the original models use miRNA sequence similarity. Dbmda1 could not be computed for HMDD3 because of out of memory error.

|  | Method | GIP |  | Functional+Semantic |  | miRNA sequence+Semantic |  |
| --- | --- | --- | --- | --- | --- | --- | --- |
|  |  | AUROC | AP | AUROC | AP | AUROC | AP |
| HMDD2 | DBMDA | 0.798 ± 0.016 | 0.022 ± 0.003 | 0.721 ± 0.007 | 0.013 ± 0.001 | - | - |
|  | DBMDA1 | 0.807 ± 0.006 | **0.025 ± 0.001** | 0.785 ± 0.005 | **0.021 ± 0.001** | - | - |
|  | DBMDA2 | 0.804 ± 0.009 | 0.024 ± 0.002 | 0.723 ± 0.007 | 0.014 ± 0.000 | 0.720 ± 0.007 | 0.013 ± 0.000 |
|  | DBMDA3 | 0.807 ± 0.007 | **0.025 ± 0.001** | 0.786 ± 0.005 | **0.021 ± 0.001** | 0.785 ± 0.007 | **0.021 ± 0.001** |
| HMDD3 | DBMDA | 0.826 ± 0.008 | 0.018 ± 0.002 | 0.757 ± 0.029 | 0.011 ± 0.002 | - | - |
|  | DBMDA2 | 0.837 ± 0.011 | 0.020 ± 0.002 | 0.783 ± 0.005 | 0.012 ± 0.000 | 0.685 ± 0.029 | 0.007 ± 0.001 |
|  | DBMDA3 | 0.839 ± 0.005 | **0.021 ± 0.001** | 0.869 ± 0.026 | **0.038 ± 0.005** | 0.801 ± 0.004 | **0.014 ± 0.000** |

**Table 7.** Mean AUROC and AP scores (with standard deviation) for Epmda and its variants on HMDD2 and HMDD3 datasets. The original model used GIP kernel similarity. Because of high time complexity, only 5 runs were performed for HMDD3 dataset. For the same reason, the scores corresponding to the last column could not be computed.

|  | Method | GIP |  | Functional+Semantic |  | miRNA sequence+Semantic |  |
| --- | --- | --- | --- | --- | --- | --- | --- |
|  |  | AUROC | AP | AUROC | AP | AUROC | AP |
| HMDD2 | EPMDA | 0.794 ± 0.040 | 0.130 ± 0.019 | 0.94 ± 0.006 | 0.382 ± 0.020 | 0.921 ± 0.004 | 0.200 ± 0.011 |
|  | EPMDA1 | 0.855 ± 0.008 | 0.170 ± 0.008 | 0.944 ± 0.006 | **0.459 ± 0.019** | 0.922 ± 0.005 | **0.216 ± 0.012** |
|  | EPMDA2 | 0.498 ± 0.015 | 0.013 ± 0.001 | 0.883 ± 0.005 | 0.144 ± 0.008 | 0.869 ± 0.004 | 0.092 ± 0.007 |
|  | EPMDA3 | 0.964 ± 0.004 | **0.504 ± 0.017** | 0.870 ± 0.009 | 0.152 ± 0.012 | 0.920 ± 0.005 | 0.155 ± 0.010 |
| HMDD3 | EPMDA | 0.798 ± 0.047 | **0.076 ± 0.007** | 0.908 ± 0.007 | 0.071 ± 0.003 | - | - |
|  | EPMDA1 | 0.870 ± 0.002 | 0.068 ± 0.002 | 0.900 ± 0.004 | **0.083 ± 0.002** | - | - |
|  | EPMDA2 | 0.493 ± 0.008 | 0.005 ± 0.000 | 0.648 ± 0.005 | 0.009 ± 0.000 | - | - |
|  | EPMDA3 | 0.775 ± 0.003 | 0.015 ± 0.000 | 0.799 ± 0.004 | 0.031 ± 0.001 | - | - |

**Fig. 3.**
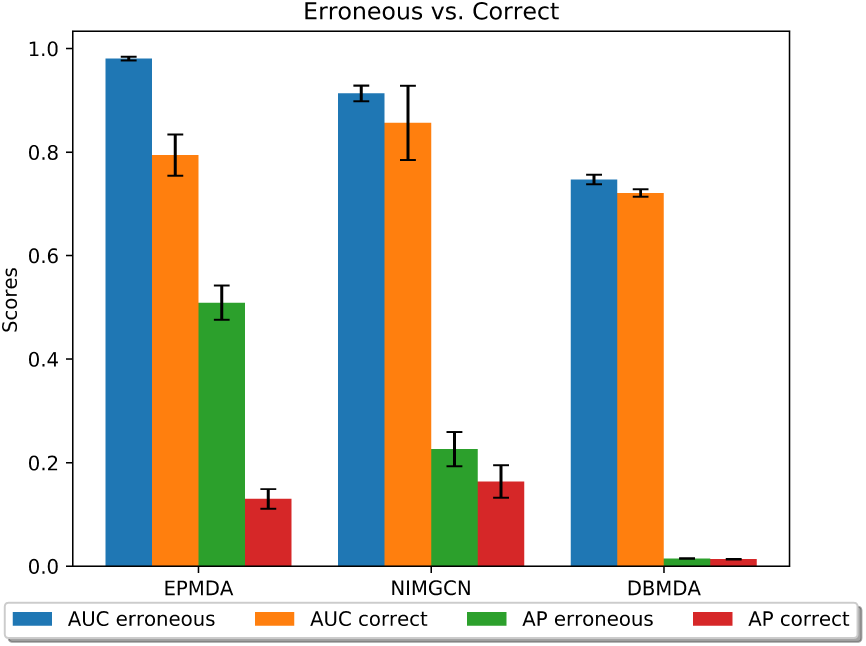
Erroneous and correct results for three original studied models on HMDD2 dataset

**Fig. 4.**
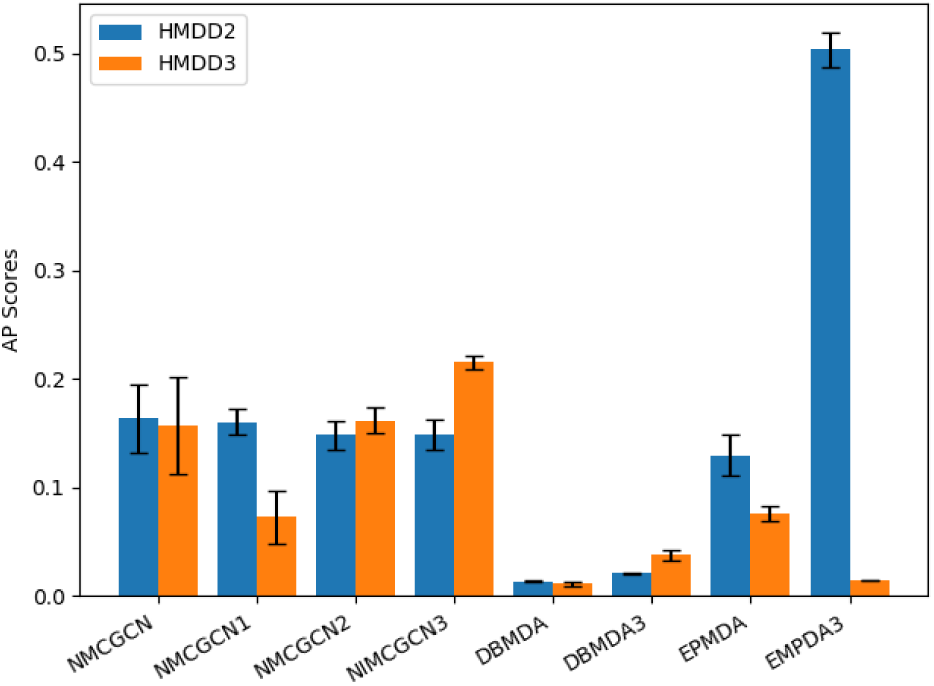
Mean AP scores (with standard deviation) for different models on HMDD2 and HMDD3 dataset. The worst performing model variants are removed from the plot. The input similarity features correspond to the ones used in the original models.

#### 4.3.2 Computation of Similarities

As already pointed out, existing works suffer from a data leakage problem in which associations from the test network were used for generating miRNA functional and `GIP` kernel similarities. In this work, we resolve the data leakage issues by re-calculating all similarities in accordance with the given data splits. More specifically, for each run, miRNA functional similarity or miRNA/disease `GIP` kernel similarity scores are computed using only the associations from the training data. That means when we do 5-fold cross validation, those similarity scores are calculated 5 times with different input.

For miRNA sequence similarity, we use the code from Dbmda. To compute `GIP` kernel similarity, we use the code provided by Epmda authors’. For the other similarity measures, we cannot get the original code to calculate the disease similarity as well as miRNA functional similarity scores given in MISIM v1.0 database. Therefore, for consistency, we use our own implementation to calculate all the miRNA functional and disease similarities (both from the whole network (as previously used by the studied models) and from only the training data alone). We provide all the code for similarities calculation at https://git.l3s.uni-hannover.de/dong/simplifying_mirna_disease.

#### 4.3.3 Hyperparameter settings

For Dbmda, since only the code for miRNA sequence similarity calculation is available, we use a simple autoencoder consisting of one encoder and one decoder. The encoder is a densely connected layer of size 32 (which is the encoded dimension explicitly given in Dbmda paper) with ReLU activation function and L1 regularization. The decoder, which has the same size as the input, is a densely connected layer with sigmoid activation function. We train that autoencoder for 1000 epoch in all experiments. For the Rotation Forest [13] classifier, we use the implementation from https://github.com/digital-idiot/RotationForest with all the default parameters. For Nimgcn, we use all the author’s parameters like the number of epochs, loss function, number of hidden units, etc. For Epmda, we use the authors’ code and settings for both feature calculation and MLP regression model.

## 5 Results and discussion

In our experiments, we investigate and answer the following research questions.

- **RQ1:** Which models are most affected by the data leakage problem caused by using precomputed similarity features?
- **RQ2:** How do our proposed model variants compare with the original models?
- **RQ3:** How sensitive are the models to different kinds of miRNA and disease similarity features (GIP, miRNA functional, disease semantic etc.)?

In Section 5.1 we answer RQ1 by comparing originally proposed models with the similarity features computed on the whole association data and using only the training split. The results corresponding to RQ2 are discussed in Section 5.2. In Section 5.3 we compare the difference in performance with respect to input similarity features (c.f. RQ3).

### 5.1 Impact of pre-computed similarities

Figure 3 shows the comparison the AUROC and AP scores of the three models on HMDD2 dataset. For all models, the “Erroneous results” denote results retrieved from models which use pre-computed similarities while “Correct results” denote results obtained from models that use similarities calculated at run time using only the given training data.

As expected, we find that the use of pre-calculated similarity does result in a much higher AUROC or AP scores. We observe the highest performance drop in Epmda because its calculated features are heavily relied on the fed `GIP` kernel similarities. We believe that the reason behind this observation is that the `GIP` similarities are computed using only the association information. To represent a disease node, Nimgcn and Dbmda use disease semantic similarity features which (unlike `GIP` based disease similarity) are computed from the disease ontology and not from the miRNA-disease associations. We, therefore, observe a lower impact of this correction step on Dbmda and Nimgcn. Moreover, Dbmda also employs additional miRNA sequence similarity features which can be computed independently from the associations which makes it relatively more robust to the changes in the miRNA functional similarity features. Therefore, Dbmda is impacted least of all models.

### 5.2 Impact of model architecture and training/test data setups

In order to answer RQ2, we compared the 3 studied models along with their several variants on HDMM2 and HDMM3 datasets. Figure 4 presents the mean AP scores for these models with standard deviation. We make the following observations.

1. Epmda3, which corresponds to our proposed variant of Epmda, outperforms all other models with a large margin(around 287.7% of improvement in AP score) for the Hmdd2 dataset. We believe that the use of balanced training data as also discussed earlier have contributed to a better trained model.
2. Nimgcn3, which is also one of our proposed variants outperforms other models for Hmdd3 dataset. Moreover, it shows a lower standard deviation over the 25 runs as compared to the original Nimgcn model.
3. From the above two observations we conclude that the edge perturbation features based on fixed cycle lengths used by Epmda are not generalizable for the larger Hmdd3 dataset. We believe that its performance can be improved by fine-tuning the hyperparameter corresponding to the length of the longest cycle.
4. Both Nimgcn and Epmda consider actively the topological structure of the heterogeneous miRNA-disease network whereas Dbmda simply extract low dimensional representations from the similarity features. The poor performance of Dbmda as compared to other models points to the importance of learning from the topological structure of the network.

Detailed results for all models and its variants using different kinds of similarity features (discussed in next section) are presented in tables 5, 6 and 7. We would also like to point out that among all the studied models, Epmda took the longest time to run. To calculate the features, we let 5 threads running in parallel. However, it still took Epmda 35 hours to run on Hmdd2 (25 runs) and nearly 27 days to run on Hmdd3 (only 5 runs) on our servers of 16 cores, 128G memory. The theoretical time complexity of Epmda is *O*(*ℓn*_*m*_*n*_*d*_(*n*_*m*_ + *n*_*d*_)^3^) where *f* is the length of longest cycles considered in graph *G* that are used to calculate the edge disturbance level, *n*_*m*_ is the number of and *n*_*d*_ is the number of diseases. Therefore, for large network, Epmda feature calculation is extremely expensive. Moreover we also note that Epmda performance on Hmdd3 is lower than Nimgcn and its variants by a large margin.

#### 5.2.1 The rate of negative training samples in the training data

As discussed earlier, the use of an imbalance training dataset might downgrade the model performance. To quantify the effect of different training data set up on the model performance, we use both the balance and imbalance training data to train our investigated models^5^. Interestingly, for Nimgcn and its variant, the imbalance training set slightly boost up those models performance but it is not the case for Hmdd3. For Epmda, it works the other way round: balance training dataset results in a much higher AP score on Hmdd2 but on Hmdd3, it produces lower scores. Since the best AP score on Hmdd2 is acquired by Epmda with balance training set and the best AP score on Hmdd3 is achieved by Nimgcn3 with balance training set, we recommend the use of balance training set.

### 5.3 Impact of input similarity features

As discussed in Section 2, the input features corresponding to miRNA and disease nodes can be computed in several ways. In particular for disease, we can compute disease semantic, phenotype and `GIP` kernel similarities. Epmda and Dbmda uses disease semantic similarity whereas Epmda uses `GIP` kernel based similarity. For miRNA, we have miRNA functional similarity (used by Dbmda and Nimgcn), `GIP` kernel based miRNA similarity (used by Epmda) and miRNA sequence similarity (used by Dbmda). In figures 6 and 5 we report mean AP scores comparing various models when they use (i) `GIP` similarity for disease and miRNA nodes (ii) `miRNA sequence+Semantic` similarity for Hmdd2 and Hmdd3 datasets respectively. We observe that `GIP` kernel based similarity measure which is directly computable on the association network and requires no other external information is in fact shows better performance in most of the models.

**Fig. 5.**
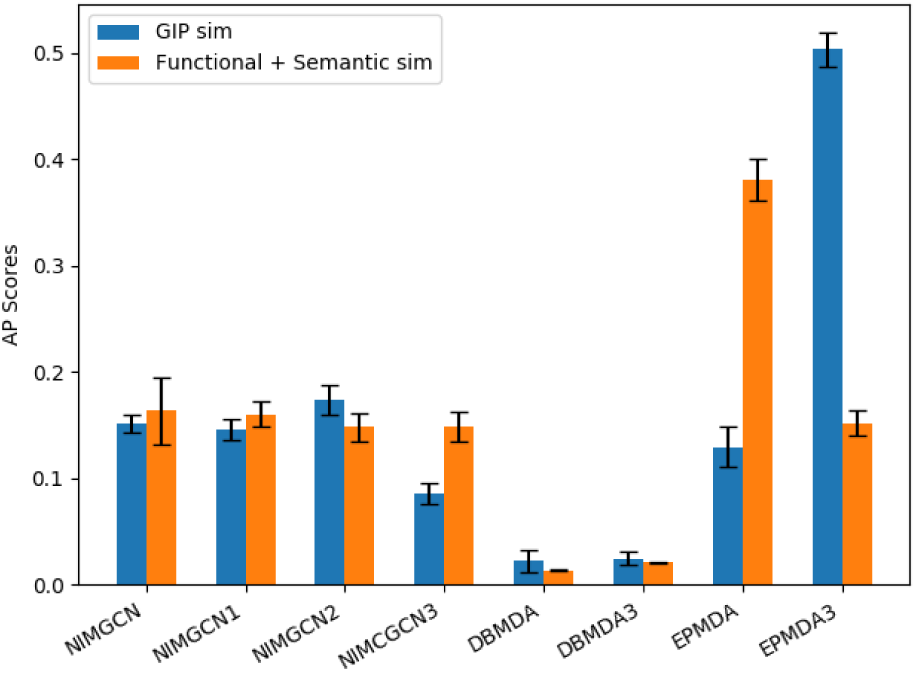
Mean AP scores (with standard deviation) corresponding to different similarity measures on HMDD2. The worst performing models are not shown in the plot.

**Fig. 6.**
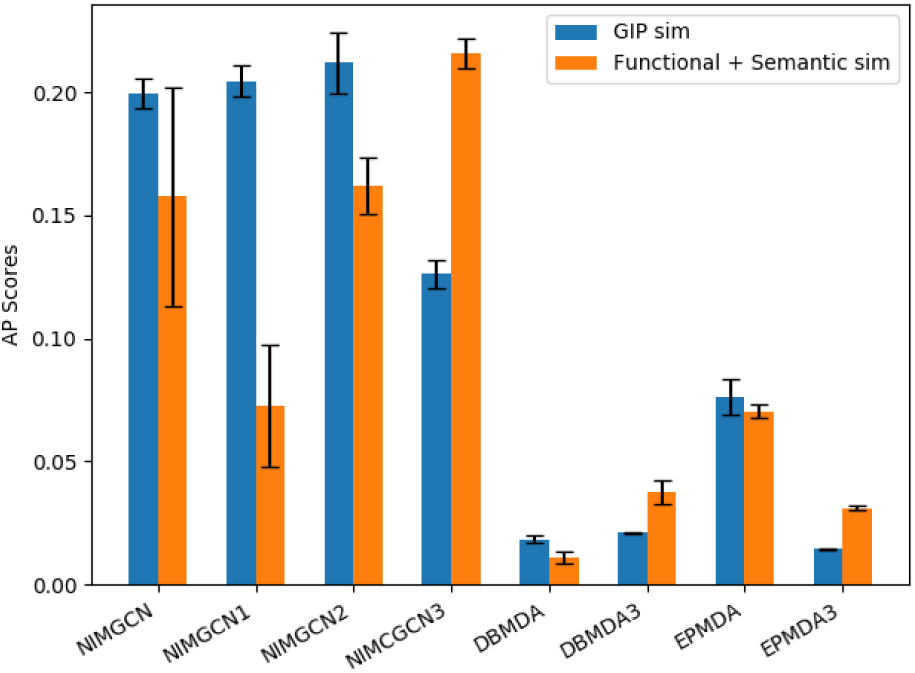
Mean AP scores (with standard deviation) corresponding to different similarity measures on HMDD3. The worst performing models are not shown in the plot.

We now discuss the detailed results presented in tables 5, 6 and 7. Note that the column `GIP` refers to the case when for both miRNA and disease `GIP` kernel similarity features are used, `Functional+Semantic` denotes that miRNA fucntional similarity and disease semantic similarity were used and `miRNA sequence+Semantic` corresponds to using miRNA sequence similarity as input miRNA features and disease semantic similarity.

#### miRNA sequence similarity

In terms of miRNA sequence similarity, we observe that in Table 6, Dbmda2 (Dbmda without miRNA sequence similarity) performs slightly better than Dbmda and Dbmda1 performs comparable to Dbmda3 (Dbmda1 without miRNA sequence similarity). That means removing miRNA sequence similarity from Dbmda or Dbmda1 does not have any significant impact on those models performance. Therefore, it is obvious that miRNA sequence similarity has very little effect on Dbmda performance.

The results corresponding to the use of miRNA functional and miRNA sequence similarity as the only source of features for miRNA are presented in the `Functional+Semantic` and `miRNA sequence+Semantic` columns of tables 5–7, respectively. Taking a closer look at the given AP scores we can see that overall, for the two experimented data sets, miRNA functional similarity which takes into account the known miRNA-disease associations is more informative than miRNA sequence similarity. However, miRNA sequence similarity still has its benefit for new miRNA where no known miRNA-disease association is available. When miRNA sequence similarity is to be used, Nimgcn2 or Nimgcn3 should be given more preference.

#### Effect of GIP kernel similarity

Figure 5 and 6 present the results for different models with different similarity measures as input. Looking at the plots we can see that on Hmdd2, `GIP` similarities, which are based on the Gaussian Interaction Profile kernel, results in better performance of Nimgcn2 (which is built up from neural projections only) but significantly lower performance of Nimgcn3 on both data sets. In addition, Epmda with `GIP` similarities result in the highest AP scores on Hmdd2, which is significantly higher than the other methods. On Hmdd3, though the highest AP score is the result of Nimgcn3 and `Functional+Semantic`, AP score of `GIP` similarities in other methods tend to be higher than the other combination.

Additionally, we also experimented with the two different disease similarity scores as described in equations (2) and (4). However, we find that overall, the two similarity measures perform comparable to each other and therefore we present results only corresponding to using disease semantic similarity given in Equation (4).

## 6 Conclusion and recommendations

In this work, we investigate three state-of-the-art models and several simpler variants of them thereby comparing a total of 12 models on a common experimental framework which we also release for reproducibility of our results and future development. We report AP and AUROC scores by running 5 times 5-Fold cross validation. For each fold, we use a complete test set miRNA-disease associations (consisting of all possible pair combination except the positive links in the training set) and recompute the similarity features each time by using the information only from the training set.

We unearth the problem of data leakage in machine learning models which employ pre-computed miRNA functional similarities or `GIP` kernel similarities computed over the complete association network. As a part or whole of test set associations were used in the feature computation, the reported results overestimate actual performance. In this work, we computed similarity features each time corresponding to the training set and showed that the Epmda model relying completely on association network for `GIP` kernel based miRNA and disease similarity features show a larger drop in performance when correct feature computation method was used.

In addition we showed that several of our proposed simpler variants of the studied state of the art techniques outperform the actual models in our experimented data sets. Our proposed variants have resulted in improvements in Average Precision scores (as compared to original models) by approximately 287.7% and 36.7% on HMDDv2.0 (AP: 0.504) and HMDDv3.0 (AP: 0.216) datasets respectively. Apart from measuring the importance of various components of the actual models, we also compare the models using different kinds of miRNA and disease similarity features. The results suggest that the use of `GIP` similarities as input often produce higher performance and hence is recommended for use in future models.

Our results indicate that a carefully chosen set of input features works better than adding more complexity/non-linearity to the model. We also find that the methods which take into account of the topological structure of the miRNA-disease network perform better. We discover that one of Epmda variants which first extracts edge perturbation based features is the best performing method for Hmdd2 dataset. For the larger Hmdd3 dataset, Epmda’s feature extraction step is very expensive and the hyperparameter corresponding to the length of longest cycle might need to be fine tuned to improve its performance on Hmdd3. We therefore recommend the design of deep learning methods based on graph neural networks for dataset dependent automatic feature extraction from the graph topology.

## 7 Acknowledgement

We would like to thank the authors of Epmda and Nimgcn for sharing their code, data and spending time in answering our questions.

## Footnotes

1 http://www.ncbi.nlm.nih.gov/mesh

2 http://www.nlm.nih.gov/

3 https://scikit-learn.org/stable/modules/generated/sklearn.metrics.auc.html##sklearn.metrics.auc

4 https://en.wikipedia.org/w/index.php?title=Information_retrieval&oldid=793358396##Average_precision

5 The scores for Nimgcn, Dbmda and their variants are not reported here as significant improvements in performance were not observed

